# BMI and BPH correlate with urinary microbiome diversity and lower urinary tract symptoms in men

**DOI:** 10.1101/2023.12.14.571758

**Authors:** Kate R. Bowie, Mark Garzotto, Eric Orwoll, Lisa Karstens

**Affiliations:** Department of Biomedical Engineering, Oregon Health & Science University, Portland, Oregon, USA; Cancer Early Detection Advanced Research (CEDAR), Knight Cancer Institute, Oregon Health & Science University, Portland, OR, USA; School of Medicine, Oregon Health & Science University, Portland, OR, USA; Portland VA Medical Center, Portland, Oregon, USA; Division of Endocrinology, Diabetes, and Clinical Nutrition, Oregon Health & Science University, Portland, Oregon, USA; Department of Medical Informatics and Clinical Epidemiology, Oregon Health & Science University, Portland, Oregon, USA; Department of Obstetrics and Gynecology, Oregon Health & Science University, Portland, Oregon, USA

**Author notes:** Correspondence to: L Karstens, PhD, Oregon Health & Science University Mailcode BICC, 3181 SW Sam Jackson Park Rd Portland, OR 97239, USA.

## Abstract

Several studies have identified bacteria and other microbes in the bladder and lower urinary tract in the absence of infection. In women, the urinary microbiome has been associated with lower urinary tract symptoms (LUTS), however, similar studies have not been undertaken in large cohorts of men. Here we examine the urinary microbiome and its association with LUTS in a subset of 500 men aged 65 to 90 years from the Osteoporotic Fractures in Men (MrOS) study. We identified significant associations between benign prostatic hyperplasia (BPH), age, and body mass index (BMI) with several diversity metrics. Our analysis revealed complex relationships between BMI, BPH, LUTS, and alpha diversity which give insight into the intricate dynamics of the urinary microbiome. By beginning to uncover the interrelationships of BPH, BMI, LUTS, and the urinary microbiome, these results can inform future study design to better understand the heterogeneity of the male urinary microbiome.

## Introduction

The bladder and urinary tract have long been considered sterile in the absence of infection; however, several studies have found evidence of a resident microbiome in both males and females.^1–3^ The urinary microbiome encompasses the bacteria, archaea, fungi, and viruses that inhabit the bladder and urinary tract, and these microbes may play a role in diseases and disorders affecting the urogenital system. The majority of urinary microbiome studies have examined the female urinary microbiome and found there are various healthy microbiome compositions commonly dominated by *Lactobacillus, Gardnerella, Streptococcus, Staphylococcus, Corynebacterium,* and *Escherichia*.^4,5^ Urinary microbiome studies in men have revealed several of the same bacteria found in women, such as *Escherichia, Lactobacillus,* and *Streptococcus*, but the male urinary microbiome can also be dominated by genera such as *Prevotella* and *Enterococcus*.^6–8^

Importantly, differences in the types of bacteria found in the bladders of women have been associated with recurrent urinary tract infections,^9^ as well as lower urinary tract symptom (LUTS) such as urge incontinence (the sudden urge to urinate followed by leakage of urine).^10,11^ The composition of the female urinary microbiome has also been found to be predictive of response to urgency urinary incontinence treatment, with a positive treatment response observed in patients with a less diverse microbiome.^9^ These studies have identified specific associations between the urinary microbiome and urological diseases, and indicate more broadly that the urinary microbiome may play a role in human health and disease.

Despite the numerous studies undertaken in women, far less is known about the urinary microbiome in men.^12^ Similar to the female urinary microbiome, there are studies demonstrating that alterations in male urinary microbiome composition have been associated with diseases and disorders. A “male urinary microbiome” literature search in 2023 resulted in 22 primary research articles. Researchers have identified potential relationships between the male urinary microbiome and kidney stones,^12^ benign prostatic hyperplasia (BPH),^13^ as well as bladder and prostate cancers,^14,15^ among many other disorders. Nonetheless, the urinary microbiome studies undertaken in men have had small cohorts (n < 100) and few have directly investigated the potential link between body mass index (BMI), age, and the male urinary microbiome. These associations could impact future study design and analysis. Larger studies are needed to both study and evaluate the broad applicability of these findings in males.

Additionally, although clear associations have been found between the female urinary microbiome and LUTS, the same associations have not been identified in a large cohort of men. Approximately one-third of men over the age of 50 experience moderate to severe LUTS.^16^ The main cause of LUTS is thought to be BPH, which is hypothesized to press on the bladder and/or urethra; however, this may not be the underlying etiology for all LUTS in all men. Here, we evaluated the urinary microbiome from a large cohort of older community-dwelling men using urine samples from the NIH-funded Osteoporotic Fractures in Men (MrOS) study ^17^. Our goals were to identify microbiome associations with clinical characteristics such as age, BMI, BPH, and general medical history, as well as identify associations with LUTS, including symptoms related to both urgency (irritation) and pushing or straining (obstruction).

## Results

### Clinical characteristics of study participants

We utilized urine samples and phenotypic data from 500 men at baseline as part of the MrOS study.^17,18^ These 500 men were randomly selected from the overall cohort of 5994 MrOS participants. The average age of the participants in the current analysis was 73.0 ± 5.7 years, with an average BMI of 27.9 ± 3.8 kg/m^2^. (**Table 1**). Participants completed a questionnaire detailing medical history, current health status, and overall quality of life. All participants provided a morning second-voided urine sample and completed a LUTS assessment using the International Prostate Symptom Score (IPSS). Just over half of participants (55.8%) had IPSS scores of 7 or lower and were designated as men with no/mild LUTS, while 44.2% of participants had scores higher than 7 and were designated as men with moderate to severe LUTS. All voided urine samples underwent 16S rRNA amplicon sequencing of the V4 region to determine urinary microbiome composition (see Methods). None of the participants reported a current urinary tract infection or recent use of antibiotics.

**Table 1.**
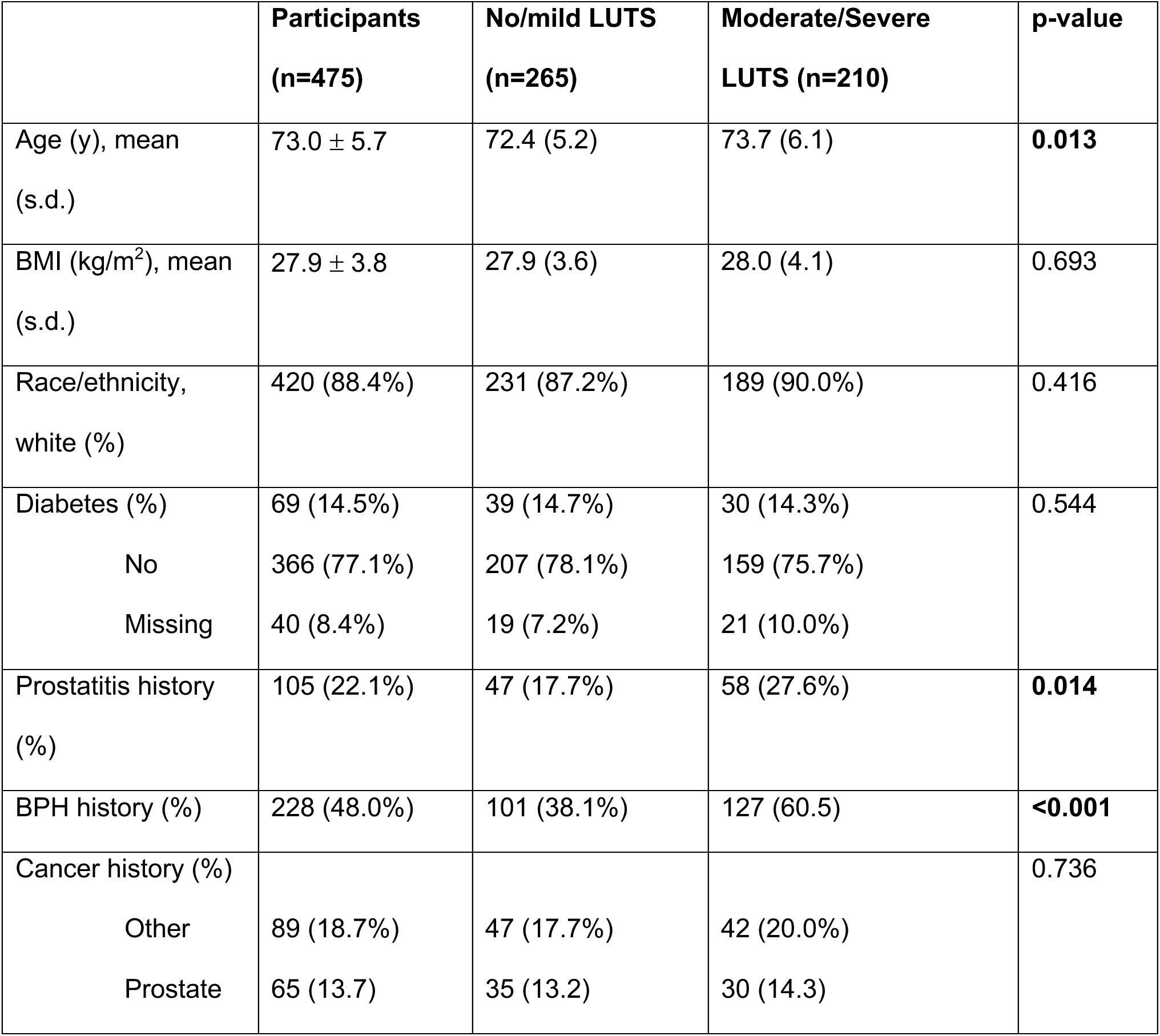
Clinical characteristics of the men from the MrOS study with urinary microbiome data.

Grouped by LUTS status, the mean age in men with moderate to severe LUTS (73.7 years) was slightly but significantly higher (n=0.013) compared with those with mild to moderate LUTS (72.4 years). There was a significant difference (p < 0.001) in the prevalence of reported BPH between the two groups, with 60.5% of men with moderate to severe LUTS having a history of BPH, compared with 38.1% in the no to mild LUTS group. Significantly more men with moderate to severe LUTS also had a history of prostatitis (27.6%) compared with those with no/mild LUTS (17.7%, p = 0.014). The men with moderate to severe LUTS having both higher rates of BPH and prostatitis is expected, as an enlarged prostate and inflammation are common risk factors for more severe LUTS.^19^ There was, however, no correlation between BMI and LUTS severity, as both groups had a similar BMI (no to mild- 27.9 kg/m^2^; moderate to severe- 28.0 kg/m^2^) in contrast to one prior study.^16^ There were also no significant differences in multiple baseline clinical characteristics, such as prevalence of diabetes, which was approximately 14% of the men in each group, or race (**Table 1**).

### BPH, BMI, and age are drivers of male urinary microbiome diversity

Of the total cohort, 25 samples were removed due to insufficient sequencing reads (<1000 reads), leaving a total of 475 samples for subsequent analysis. Like other body sites, the male urinary microbiome composition varied between individuals and is highly heterogenous (**Figure S1a, Figure S2**). Twenty-one phyla and 571 genera were identified in the male urinary microbiome, with 54 genera being core members (present in at least 10% of samples).

Firmicutes was the most abundant phyla with a mean of 40.2% ± 25.7%, followed by Proteobacteria, Actinobacteria, and Bacteroidetes (27.4% ± 27.9%, 15.0% ± 15.4%, and 13.4% ± 15.7%, respectively **Figure S1b**). In the majority of samples, either Firmicutes or Proteobacteria was the dominant phyla, consistent with prior male urinary microbiome studies.^1,20,21^ The five most abundant genera identified were *Staphylococcus*, *Neisseria*, *Corynebacterium*, *Prevotella*, and *Streptococcus* (**Figure S1c**).^1,21^

We next evaluated if the diversity of the male urinary microbiome was associated with specific clinical characteristics (**Table Sup2, Figure 1a**). We identified significant associations between BPH and several measures of alpha diversity of the urinary microbiome (Shannon, Inverse Simpson, and Pielou indices, p < 0.05). We found that men with BPH tended to have increased alpha diversity when compared with men without BPH. We did not identify any significant associations between BPH and any beta diversity measures (weighted UNIFRAC, Unweighted UNIFRAC, Bray-Curtis), but all three beta diversity measures were significantly associated with both BMI and age (**Figure 1a**). We also did not identify any significant associations with urinary alpha diversity and age, BMI, diabetes, prostatitis, or race.

**Figure 1.**
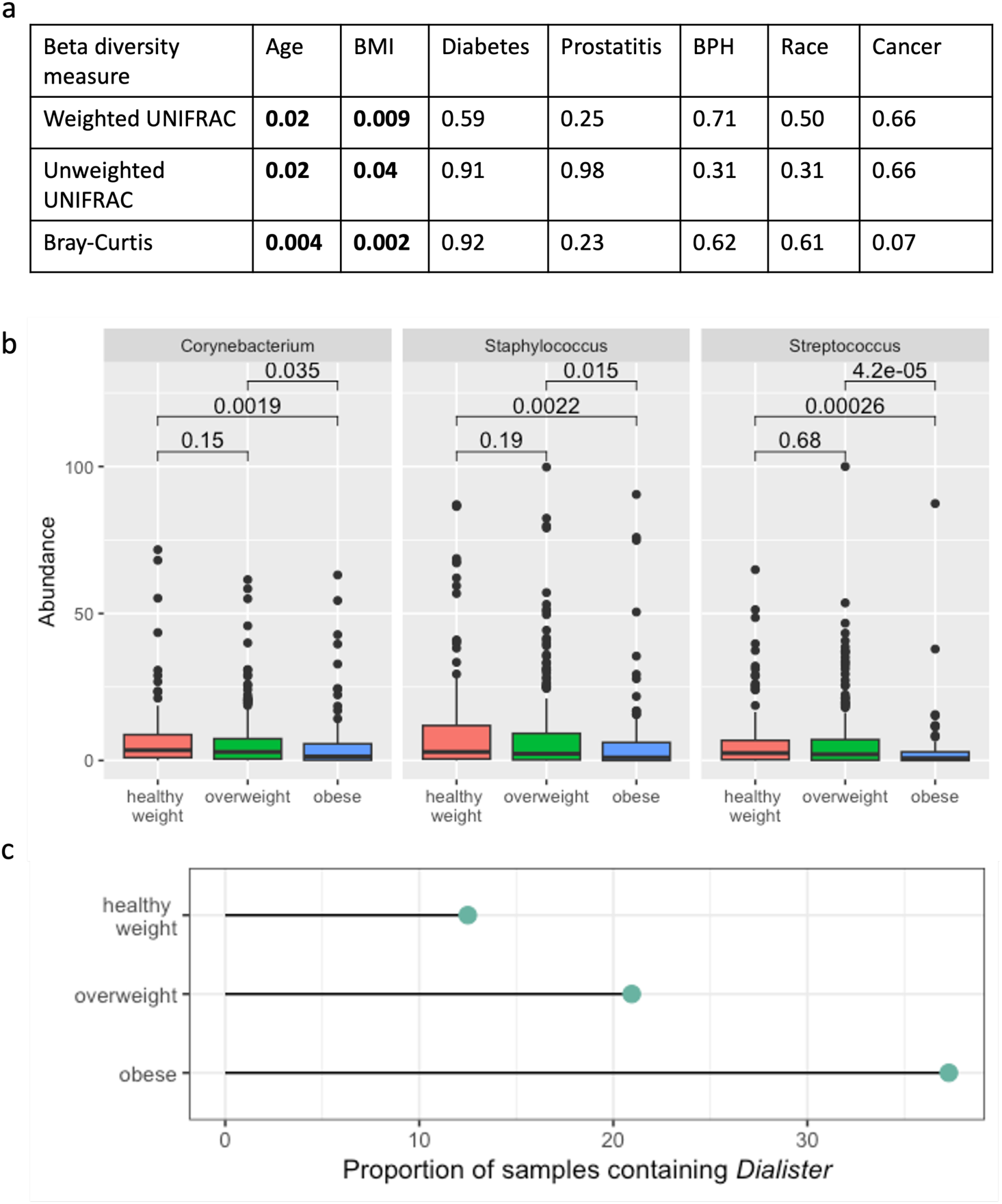
BMI is a dominant influence on beta diversity in this population. **a**) Table of p-values of PERMANOVA testing the relationship of the clinical characteristics with the beta diversity metrics of the urinary microbiome. BMI and age are both significantly associated with all three beta diversity metrics (bolded). **b**) Boxplots of relative abundance of *Corynebacterium, Staphylococcus,* and *Streptococcus* for men at healthy weight (red), overweight (green), and obese (blue) BMIs. **c**) Lollipop plot showing the percent of samples that contain *Dialister* by BMI grouping. A significantly higher proportion of obese individuals have *Dialister* in their urinary microbiomes as compared with both overweight and healthy weight individuals (p < 0. 00001).

Based on the associations between BMI and beta diversity, we next compared the relative abundances of the five most abundant phyla and 10 most abundant genera between men with BMIs in the healthy, overweight, and obese ranges. We found no significant differences in relative abundances at the phylum level. However, we identified differences at the genus level for three genera (*Corynebacterium, Staphylococcus,* and *Streptococcus*, (**Figure 1b, Figure S3**). Overall, obese men had the lowest abundance of *Corynebacterium, Staphylococcus,* and *Streptococcus*, whereas healthy men had the highest relative abundances. All three genera are known members of a healthy male urinary microbiome,^1,8^ and significant differences in relative abundances of these taxa have been described in the gut microbiomes between obese individuals and those with lower BMIs.^26^ We also examined the 5 most abundant phyla and 10 most abundant genera between individuals with and without BPH and of different age groups (determined by tertile). We found no significant differences with BPH (data not shown); however, the urinary microbiomes of men 76 to 90-years old had significantly higher relative abundance of Actinobacteria compared to men 71 to 75-years-old (p = 0.03, **Figure S4**).

We next determined how specific taxa in the male urinary microbiome were associated with age, BMI, BPH, and the remaining clinical characteristics reported in **Table 1**. To accomplish this, we employed an exploratory analysis using the machine-learning method, Hierarchical All-against-All (HAllA) association testing, which is high-sensitivity pattern discovery in large, paired multi-omic datasets.^27^ HAllA detects possible associations with specific taxa and accompanying metadata using a statistical method for discovery. After applying HAllA at the phylum, family, and genus levels, we discovered a significant association between BMI and the genus *Dialister* (p = 0.026). *Dialister* has previously been reported in both the female and male urinary microbiome, as well as in the gut microbiome.^6,28^ In the gut, higher abundances of *Dialister* were found in people with a higher BMI and is associated with difficulty in losing weight.^29–31^ Similarly, in our data we found *Dialister* to be present in 23.2% of samples (**Figure 1c**), with the highest proportion in obese men, followed be overweight men, and lastly healthy weight men (37.3%, 20.9%, and 12.5% respectively, p<0.0001, **Figure 1c**). We also identified an association between the prevalence of diabetes and the phylum Bacteroidetes (p = 0.017), which has been shown to have a similar association in the gut microbiome, but has not been previously described in the male urinary microbiome.^32^ We did not identify associations between specific taxa and age or BPH using HAllA. Although age and BPH are significantly associated with beta diversity and alpha diversity, respectively, these HAllA results do not point to any specific bacteria. This suggests that multiple bacteria in the microbiome community likely contribute to the differences seen above in diversity.

### Clustering reveals 8 “Urotypes”

After examining specific taxa and their associations with clinical characteristics of this MrOS cohort, we sought patterns in the overall urinary microbiome composition. In view of the associations between the urinary microbiome and BPH, BMI, and age, we determined if the associations between BMI, age, and BPH vary in the context of different microbiome compositions. We used Dirichlet multinomial modeling (DMM) to cluster samples with similar urinary microbiome compositions into “urotypes.”^33^ DMM clustering on members of the core urobiome revealed eight urotypes each dominated by a specific bacterium (**Figure 2a**). Each urotype contains between 19 and 89 samples. Urotypes 1, 2, and 7 were dominated by *Staphyloccocus*, while the rest were dominated by *Corynebacterium*, *Neisseria*, *Prevotella*, and *Anaerococcus*. Many clusters are dominated by a single bacterium while others are more diverse – for example cluster 5 is predominantly *Neisseria*, while cluster 4 is more diverse with high proportions of *Neisseria*, *Streptococcus*, and *Escherichia*.

**Figure 2.**
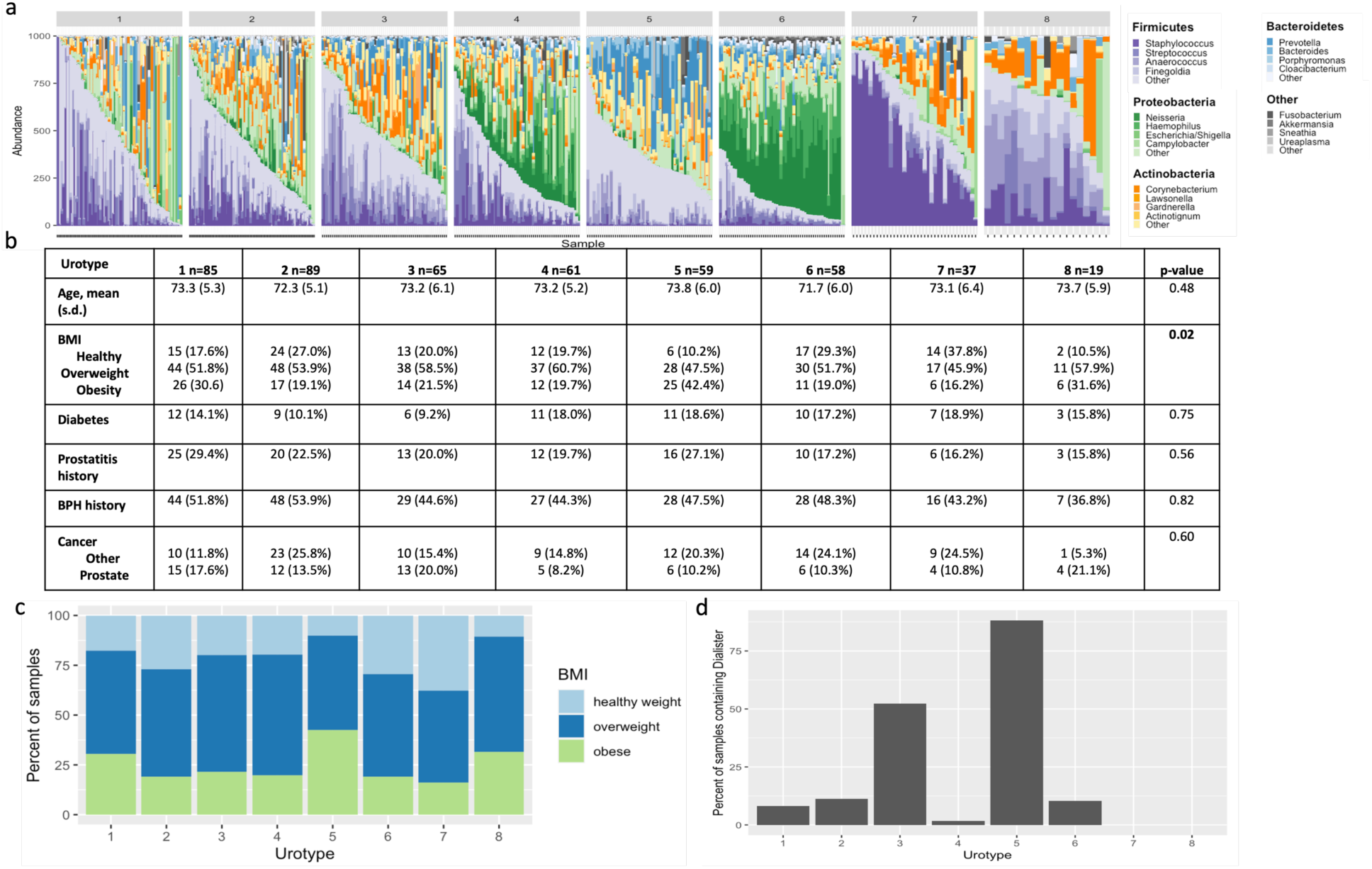
Distinct patterns of the male microbiome are present and are associated with BMI. **A)** Stacked bar plots of the eight urotypes created using Dirichlet Multinomial modeling. Each urotype is dominated by specific bacteria **b.)** Table summarizing the clinical characteristics for each urotype. BMI is significantly different between urotypes (p = 0.02, bolded) **c**) Traces of the percent of samples in each urotype colored by BMI group. **D.**) Bar graph of the percent of *Dialister* in each urotype. Urotype 5 has the highest percentage of *Dialister* and obese men.

Next, we investigated clinical characteristics within the urotypes, and observed no significant differences in age, diabetes, prostatitis, BPH, or cancer. However, in line with our previous findings which established BMI is correlated with the composition of the male urinary microbiome, we determined the urotypes had significantly different proportions of samples belonging to healthy weight, overweight, and obese individuals (p = 0.02, **Figure 2c**). Earlier we determined a relationship between *Dialister* and the urinary microbiomes of obese men; thus, we wanted to explore that connection within these urotypes. We discovered that 88.1% of the individuals of urotype 5, which has the highest percentage of obese men (42.4%) and lowest percentage of healthy weight men (10.2%, **Figure 2c**), had *Dialister* in their urinary microbiomes (**Figure 2d**). Interestingly, urotype 8 has the second most obese men (31.6%), yet none of those men had *Dialister* in their urinary microbiomes.

### BMI, BPH and LUTS characteristics are associated with the diversity of the male urinary microbiome

Next, we wanted to determine if there was an association between LUTS and the male urinary microbiome. Since our results demonstrated that BPH, BMI, and age are associated with the overall male urinary microbiome composition, and previous studies have shown that BPH and BMI can also impact LUTS, we incorporated these variables into subsequent analysis.

To evaluate whether there are associations between the microbiome and LUTS, we first looked at the complexity of the samples. Men with no to mild LUTS had an average of 16,369 reads per sample (minimum 1,018; maximum 101,017), while men with moderate to severe LUTS averaged 19,367 reads per sample (minimum 1,184; maximum 81,167; p = 0.04, **Figure S5**).

We then inspected the urinary microbiome compositions in men with no to mild LUTS compared with those with moderate to severe LUTS and did not find significant differences (**Figure 3a**). We also examined alpha diversity metrics (Kruskal-Wallis test) and beta diversity metrics (PERMANOVA analysis) of the urinary microbiome composition in these two groups, and found no significant differences were found even when adjusted for BPH, BMI, and age (**Figure 3c and Figure 3d**).

**Figure 3.**
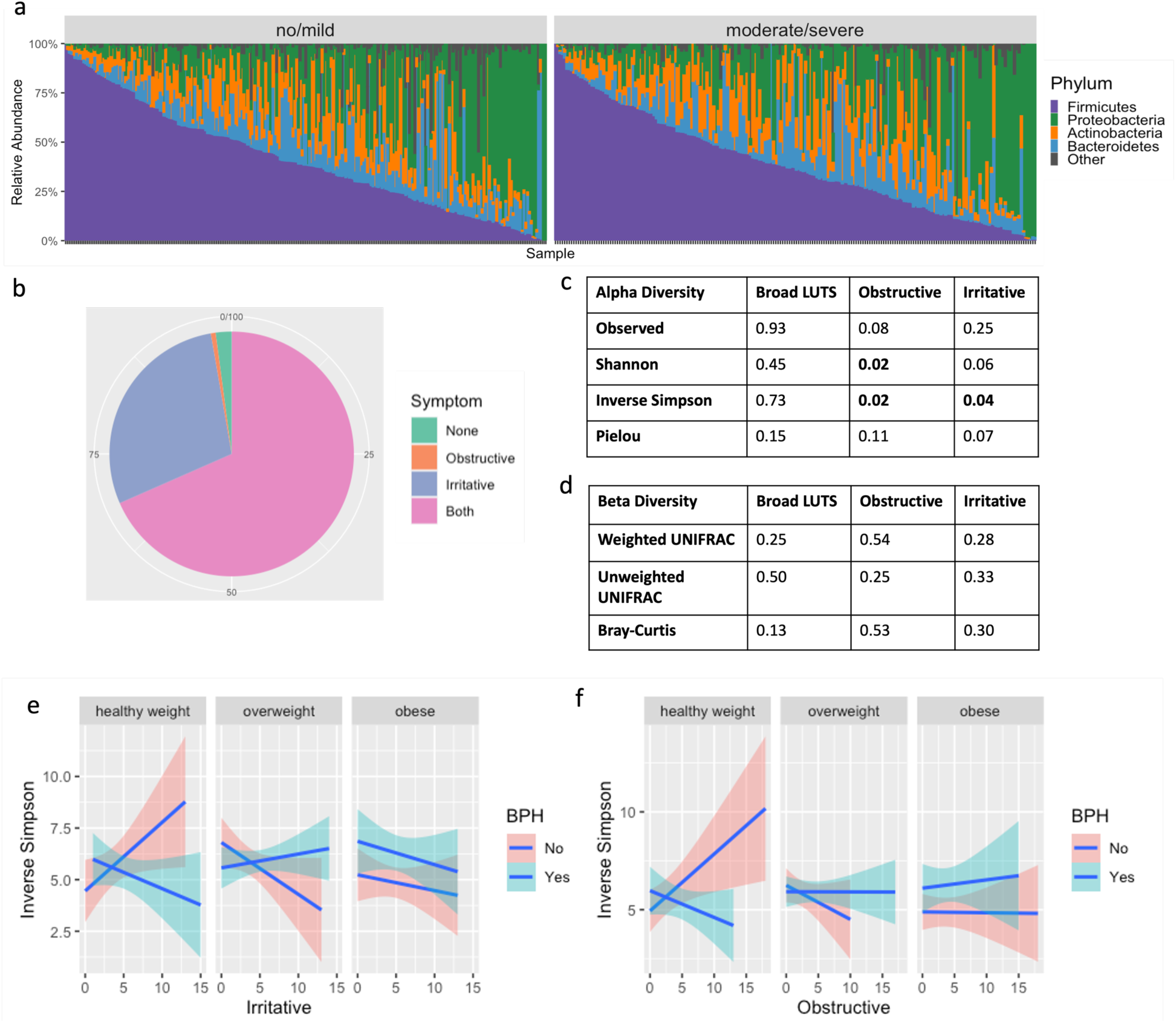
The alpha diversity of the male urinary microbiome is associated with irritative and obstructive LUTS when adjusted for BMI and BPH. **A)** Stacked bar plot of the male urinary microbiome subset by LUTS status. **B)** Pie chart showing percentage of men in the MrOS cohort who experience no symptoms, obstructive symptoms only, irritative symptoms only, and both obstructive and irritative symptoms. **C)** P-values for alpha diversity metrics and **d)** beta diversity metrics for broad LUTS (no to mild LUTS versus moderate to severe LUTS) and obstructive and irritative symptoms. Adjusted model includes interaction terms for BMI and BPH **e)** Significant interactions between BMI, BPH, and irritative or **f)** obstructive symptoms influence inverse Simpson index.

LUTS is defined broadly, encompassing both irritative and obstructive symptoms, and we hypothesized that there may be differences in the microbiome that were related to these characteristics rather than overall severity. For example, studies on LUTS and the urinary microbiome in women have found significant differences in women with stress incontinence versus urgency incontinence;^25^ however, differences are not typically found when examining overall incontinence.^34^ After separating the IPSS questions into those addressing irritative versus obstructive symptoms, we found the majority of men experienced some degree of both symptoms. Fewer had irritative symptoms alone, while less than 1% of men experienced only obstructive symptoms (**Figure 3b**). Irritative symptom scores ranged from 0 to 15, while obstructive symptoms ranged from 0 to 18.

Taking a closer look at how obstructive and irritative symptoms are associated with alpha and beta diversity, we used a Kruskal-Wallis Test and adjusted for BPH, BMI, and age. We found the Inverse Simpson alpha diversity measures significantly associated with obstructive symptoms and irritative symptoms independently (**Figure 3f**). Obstructive symptoms were also associated with the Shannon index, while the association with irritative symptoms was weaker, and not significant (Shannon, p = 0.06). In PERMANOVA comparisons, we found no associations with beta diversity and obstructive nor irritative symptoms (**Figure 3d**). In addition to the alpha diversity, we also discovered significant interactions between irritative symptoms and BPH and BMI (**Figure 3e**), as well as similar results with obstructive symptoms (**Figure 3f**). In other words, men with BPH have varying associations between alpha diversity and symptom severity depending on BMI, and there are different associations if the individual does not have BPH. For example, in the healthy weight population, men with a higher Inverse Simpson index tended to have increased irritative symptoms if they do not have BPH (**Figure 3e**). However, the converse is true if they have BPH. We see the opposite trends for men in the overweight category and found that BPH does not have the same impact in the obese group.

### Enterococcaceae and Caulobacteraceae are associated with irritative symptoms

In light of irritative and obstructive symptoms and their potential role in the diversity of the male urinary microbiome, we re-examined the eight urotypes and HAllA. There were no significant differences in the obstructive and irritative symptom scores between urotypes (p>0.05), nor when investigating broad LUTS severity.

To discover relationships with specific taxa and LUTS severity or irritative and obstructive symptoms in the older male urinary microbiome, we again employed (HAllA)^27^. Supporting our previous results, no significant associations at any taxonomic resolution were identified with overall LUTS score. We investigated irritative versus obstructive symptom scores independently, which resulted in associations at the family level, between irritative symptoms and the presence of Enterococcaceae (p = 0.002) and Caulobacteraceae (p < 0.001) independently. Enterococcaceae has previously been described in the female urinary microbiome of women with mild and moderate to severe LUTS.^35^ Caulobacteraceae has been found in the healthy canine urinary microbiome,^36^ as well as the urinary microbiome of spontaneously tolerant kidney-transplant recipients;^37^ however, no research has shown any relationship to urgency or LUTS.

We further assessed each of the questions from the American Urological Association symptom index (AUA-SI) questionnaire for a more granular analysis and found an association at the genus level between the urge to urinate frequently and *Mobiluncus* (p < 0.01). *Mobiluncus* is primarily found in post-menopausal women and women with bacterial vaginosis and has not previously been reported to persist in the male urinary microbiome.^38,39^

## Discussion

Our analysis of a relatively large cohort of older community-dwelling men revealed that the male urinary microbiome is heterogeneous and exhibits a great deal of inter-individuality in the present microbes. We found that the male urinary microbiome is dominated by *Staphylococcus, Corynebacterium, Prevotella,* and *Finegoldia*, which supports previous research (**Figure S1c**).^1,40^ We also identified complex relationships between the male urinary microbiome and BMI, BPH, age, and LUTS. While previous studies have found age-specific compositional differences in the urinary microbiome regardless of gender,^2^ associations between BMI and the urinary microbiome have been established only in women.^25^

A major finding of our study is the significant association between BMI and the male urinary microbiome composition (**Figure 1a**). Similar associations have been identified in other microbiome communities in the human body^41^ and in the urinary microbiome in women, ^25^ but have not yet been reported in the male urinary microbiome. This means that men with similar BMIs have similar urinary microbiome compositions, compared to men with different BMIs regardless of age, BPH, and medical history.

We found that BMI is linked to specific genera as well as to the overall microbiome composition. Using a Kruskal Wallis rank sum test, the relative abundances of *Corynebacterium*, *Staphylococcus*, and *Streptococcus* were significantly different in healthy weight, overweight, and obese men; obese men had a decreased abundance of all three genera, whereas healthy weight men had an increased abundance (**Figure 1b**). Since these genera are typically found in a healthy urinary microbiome, and BMI is significantly associated with beta diversity, we can hypothesize that the lower relative abundance of these three genera allows other bacteria, like *Dialister*, to create a niche in those communities. A machine-learning approach, HAllA, revealed a significant relationship between the abundance of *Dialister* and higher BMI (**Figure 1c**), an association which has not been previously reported in the context of the urinary microbiome.^29–31^ In the gut, many studies have shown *Dialister* in individuals with higher BMIs and who have difficulty losing weight.^26,30,41,42^ One study surmised that *Dialister* aggravates the host inflammatory response and insulin resistance by releasing more lipopolysaccharides,^30^ which are known to be an important feature in metabolic disease and weight gain.^43^ Interestingly, in the DMM clustering that revealed eight urotypes (**Figure 2**), urotype 5 contained the highest percentage of obese men and the highest percentage of men with *Dialister*. Although this mechanism has not been studied in the urinary microbiome, a similar one could be at play. Interestingly, in the DMM clustering that revealed eight urotypes (**Figure 2**), in the second highest cluster of obese men (urotype 7) *Dialister* was not detected. We believe this is something to investigate further and examine if other bacterial communities in the urinary microbiome could have an association with BMI.

We also observed that men with BPH tended to have a higher alpha diversity in their urinary microbiome as compared to men without (**Table Sup2**) – a finding that had previously not been reported in studies.^23^ BPH has previously been associated with significant changes in the urinary microbiota and in the microbes of the prostate tissue itself.^22,23^ In rats, BPH has been associated with the beta diversity of the gut microbiome; however, no such studies have been undertaken in humans.^24^

We did not identify significant associations between overall LUTS score and the urinary microbiome composition in men. Previous work has shown an association between more severe LUTS and the following genera: *Haemophilus*, *Staphylococcus, Dolosigranulum, Listeria, Phascolarctobacterium, Enhydrobacter, Ruminococcus, Bacillus, Faecalibacterium, and Finegoldia*.^23^ We did not find *Listeria* nor *Ruminococcus* in our dataset, and there were no associations between the other genera and LUTS severity. However, we believe the overall IPSS severity score may not allow for the detection of associations with specific causes of urinary symptoms (e.g., irritative and obstructive symptoms), which could have different etiologies. We considered the hypothesis that individual elements of the LUTS symptoms score could have different associations with the microbiome. For instance, irritative symptoms could have more inflammatory bacteria, while obstructive symptoms could be the result of bacteria more likely to create a biofilm which could create some blockage while voiding. In support of this idea, we did find significant associations between alpha diversity metrics and specific obstructive and irritative LUTS symptoms when the analyses were adjusted for BMI, BPH, and age. The inverse Simpson index was significantly associated with both obstructive and irritative symptoms; however, different trends were observed depending on BMI grouping and BPH (**Figure 3e-f**). Similar to what was discussed earlier with *Dialister* and BMI, these results suggest that there could be different mechanisms causing irritative or obstructive symptoms in obese men as compared to overweight men with or without BPH.

HAllA unveiled associations at the family level between Enterococcaceae and Caulobacteracaea with irritative symptoms; however, no associations were found at the phylum or genus level, nor any associations with obstructive symptoms. The HAllA analysis was restricted to 54 of the 571 genera, so this may have contributed to our limited findings. More work is needed to determine which taxa are important for irritative and obstructive symptoms; however, the heterogeneity of bacteria between individuals and other characteristics of the microbiome may not yield consensus. In other words, biological functions may be more homogenous and may be more indicative of phenotype or symptoms, despite being caused by a number of different bacteria.

The large sample size of this study allowed us to investigate broad trends across the urinary microbiome of older community-dwelling men. Although we had a rich dataset, there were some limitations that may have affected our analysis. Elements of the phenotypic data (e.g., history of diabetes and cancer) were self-reported and thus subject to errors. The samples collected were voided urine, which have been reported to not represent only the male bladder microbiome, but instead be a mixture of the bladder and urethral communities.^21^ Also, as with any 16S rRNA sequencing study, analysis was limited to the genus level, and resolution to the species level may improve findings. For example, there are many different species of *Dialister*. Finally, the LUTS score is based on a set of symptoms that may be the result of multiple biological pathways, and thus difficult to disentangle.

In summary, the male urinary microbiome is complex and challenging to investigate. There is considerable individual variation in the urinary microbiome in men, and that variation may be related to the presence and character of LUTS. Our study indicates that considering the specific type of LUTS is important in understanding the contribution of urinary microbiota to urinary disorders in men. We also revealed there is an intricate connection between BMI, BPH, urinary microbiome diversity, and severity of irritative and obstructive symptoms. More studies are needed to further investigate these covariates, and we believe future urinary microbiome studies in men should heavily consider BMI and BPH in study designs and analyses.

## Methods

### Study Population

We acquired samples and data from 500 randomly selected participants from the prospective NIH-funded MrOS study. The MrOS study recruited a total of 5,994 men between the ages of 65 and 100 years from 6 clinical sites in the United States to assess risk factors for fracture and other conditions related to aging. The cohort and recruitment methods have been previously described.^17^ At baseline, participants were at least 65 years of age, able to consent, walked without assistance of another person, and did not have bi-lateral hip replacement or any condition that in the judgment of the site investigator would likely impair participation in the study.^44^ The Institutional Review Boards at all sites reviewed and approved the study, and all participants provided written informed consent. Enrolled participants completed a series of medical questionnaires, including medical history of prostatitis and BPH and the AUA-SI questionnaire, and provided specimens such as blood and urine which were banked for future research. Participants were recruited between 2000 and 2002 and followed longitudinally. Morning, second-voided urine specimens were collected in sterile containers and frozen at −80 °C for future analyses.

### DNA isolation and Sequencing

Bacterial DNA was isolated from 4-mL aliquots of urine specimens. The V4 region of the 16S rRNA gene was selectively amplified to evaluate microbial composition of the urine. DNA from the 500 urine samples was submitted to Baylor University for paired-end 250 base-pair sequencing using Illumina MiSeq by Dr. Nadim Ajami.

### Classification of LUTS

LUTS was assessed using the IPSS. The IPSS is a seven-question examination that evaluates both irritative and obstructive aspects of LUTS. Each question is given a score from zero to five in terms of how severe that specific symptom is, and the scores for all questions are totaled. No/mild LUTS were assigned to scores 0 ≤ 7, while moderate/severe LUTS corresponds to scores greater than 7. LUTS was then divided into irritative versus obstructive symptoms for sub-analyses based upon the IPSS questionnaire. Scores from questions 1, 3, 4, and 6 were totaled for an overall obstructive symptom score, and the scores from the remaining questions made up the irritative score.

### Bioinformatics and Statistical Analyses

Raw sequences were processed into amplicon sequence variants (ASVs) using DADA2. The RDP Classifier was used to map the ASVs to the SILVA 128 16S rRNA reference set for taxonomic identification. The 500 samples were rarefied to 1000 reads, which removed 25 samples and left 475 samples for downstream analyses. All subsequent analyses were done on ASVs agglomerated at the genus level in R, except for application of HAllA which uses Python.^27^ Data were further processed using phyloseq (version 1.42.0) and visualized using *microshades* (version 1.10).^45^ The *Vegan* R package version 2.6.4 and *rstatix* version 0.7.2 were used for all statistical analyses.

### Testing for associations between taxa and clinical characteristics

We implemented HAllA version 0.8.20 as an exploratory analysis to investigate associations between individual taxa and the clinical characteristics. HAllA was carried out separately at the genus, family, and phylum levels, and we exported both our metadata and ASV tables to text files. The metadata included the clinical variables of Table 1, and separate ASV tables were generated for each taxonomic level. We also looked at relative abundance of the top 5 phyla and top 10 genera for BMI, BPH, and age. A Kruskal-Wallis test was used to test for a significant association between relative abundance and clinical characteristic of interest, and then a pairwise Wilcoxon Rank Sum test with false discovery rate correction.

### Clustering of samples by microbiome composition

Treating each urinary microbiome as a community, we decided to cluster based on these communities and then look for associations with clinical characteristics and LUTS. We used the *DirichletMultinomial* version 1.40.0 for DMM clustering on the “core” taxa, or the bacteria present in at least 20% of samples, to improve processing speed. We evaluated the model fit for two through 12 clusters using the Laplace approximation and chose eight clusters based on a global minimum (**Figure S5**)^33^ and found that qualitatively eight clusters exemplified eight different community types dominated by different taxa (Figure 2a). The samples were separated into eight clusters based on the core taxa, and the bacterial communities in each cluster were tested for associations with clinical characteristics using the tableone R package (version 0.13.2).

## Author Contributions

E.O. designed the study and assisted with data access and acquisition. M.G. guided data analysis and interpretation with clinical input. L.K. designed and guided the data analysis, and wrote the manuscript. K.B. analyzed the data and wrote the manuscript. All authors contributed to interpretation of results and editing the manuscript.

## Acknowledgements

The authors would like to thank members of the Osteoporotic Fractures in Men (MrOS) consortium, particularly Lynn Marshall. The MrOS Study is supported by National Institutes of Health funding. The National Institute on Aging (NIA) and the National Center for Advancing Translational Sciences (NCATS) provide support under the following award numbers: R01 AG066671 and UL1 TR000128. This project was additionally supported by NIH/NIDDK K01DK116706 (LK) and the Cancer Early Detection Advanced Research Center at Oregon Health & Science University’s Knight Cancer Institute (KB).

