## Supplementary Data for "BMI and BPH correlate with urinary microbiome diversity and lower urinary tract symptoms in men"

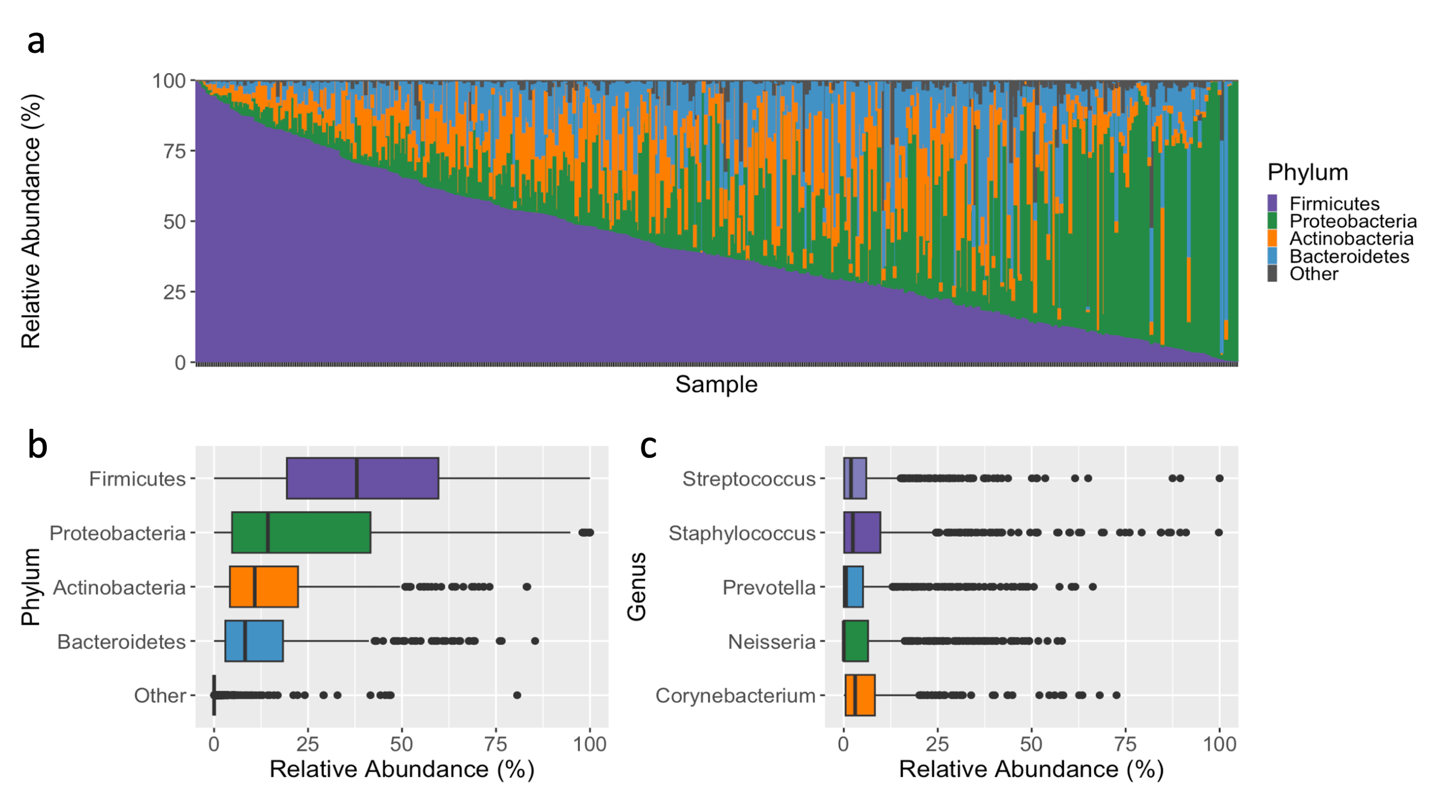

**Figure S1**. The male urinary microbiome is highly heterogenous. **a**) Stacked bar plot of the male urinary microbiome at the Phylum level ordered by Firmicutes abundance. **b**) Boxplots of the most abundant Phyla and **c**) Genera in the male urinary microbiome.

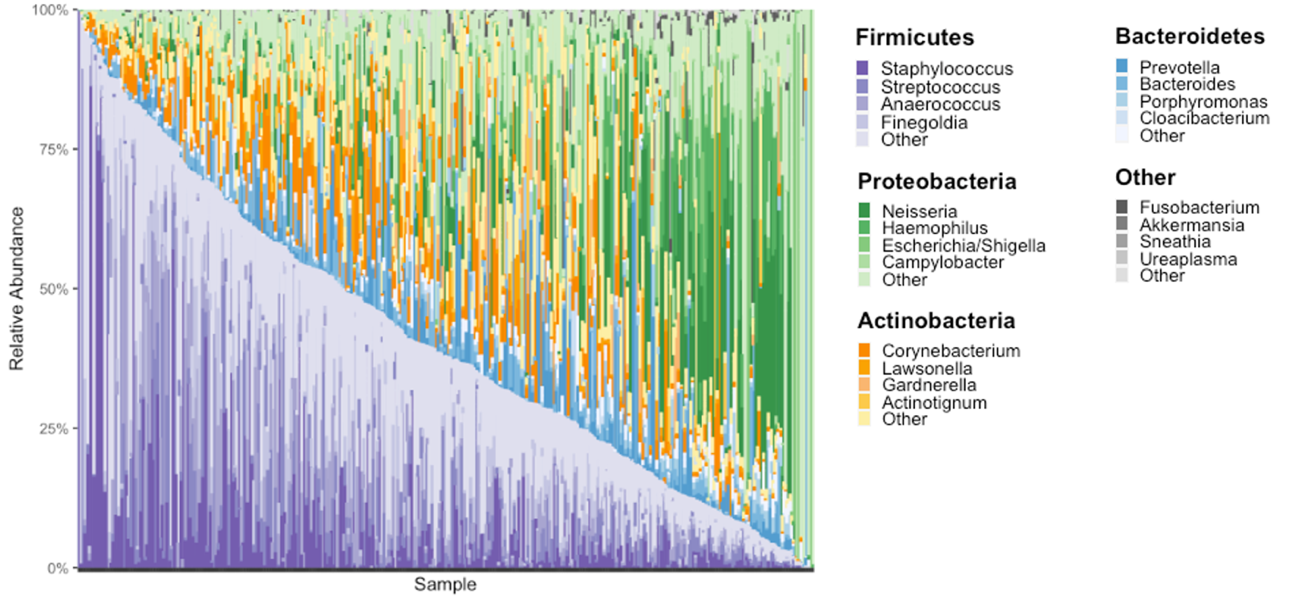

**Figure S2.** The male urinary microbiome is highly variable demonstrated by abundance of different genera. A stacked bar plot of the male urinary microbiome organized by Firmicutes abundance. Each phylum is assigned a color, and within each phylum the darker shades are assigned to the most abundant genera. The lightest shade of each color is the rest of the genera corresponding to that genera. The overabundance of light shades (i.e. in Firmicutes and Proteobacteria) demonstrates that the male urinary microbiome is highly heterogenous.

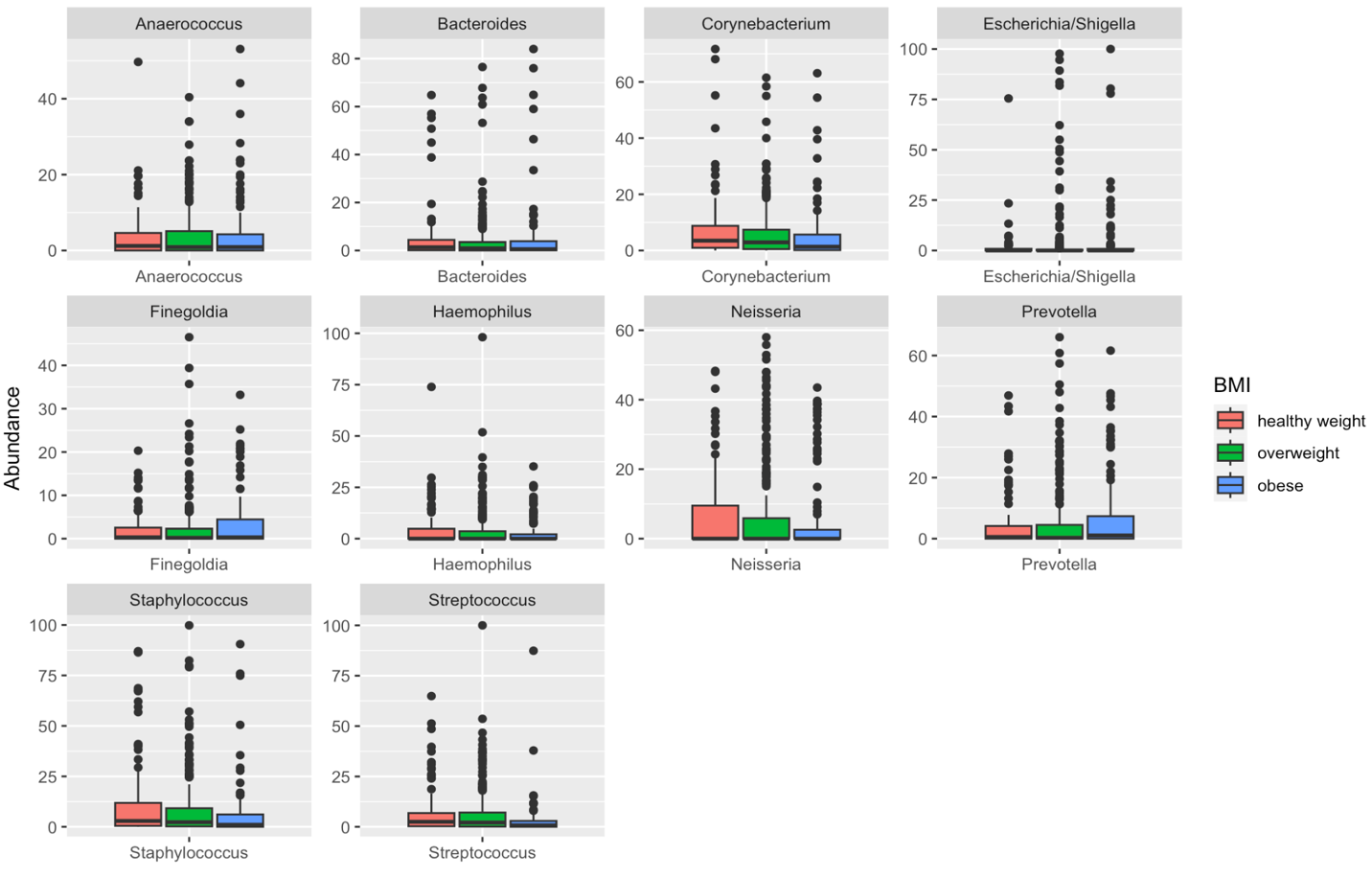

**Figure S3**. Boxplots of the most abundant genera in the MrOS cohort by BMI group (healthy weight, overweight, and obese). There were significant differences in abundance between groups detected for *Corynebacterium*, *Staphylococcus*, and *Streptococcus* highlighted in **Figure 2b**.

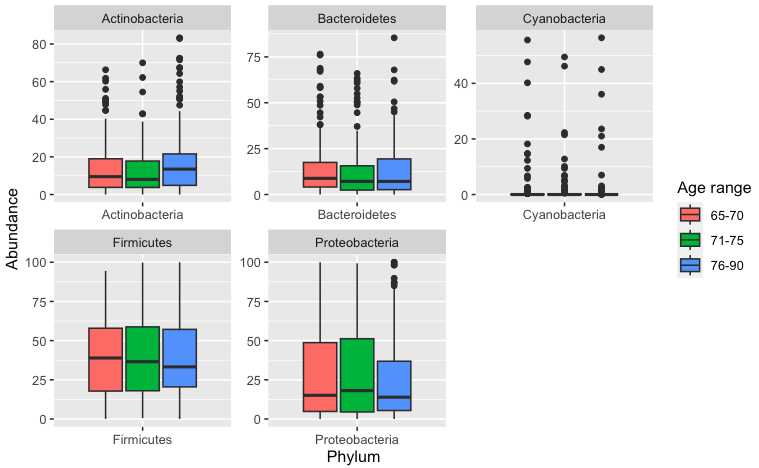

**Figure S4**. Boxplots of relative abundances of the top five most abundance phylum by age range. Age ranges were determined by tertiles. The relative abundance of Actinobacteria in men aged 76 to 90 is significantly higher than in men aged 71 to 75 (p = 0.03).

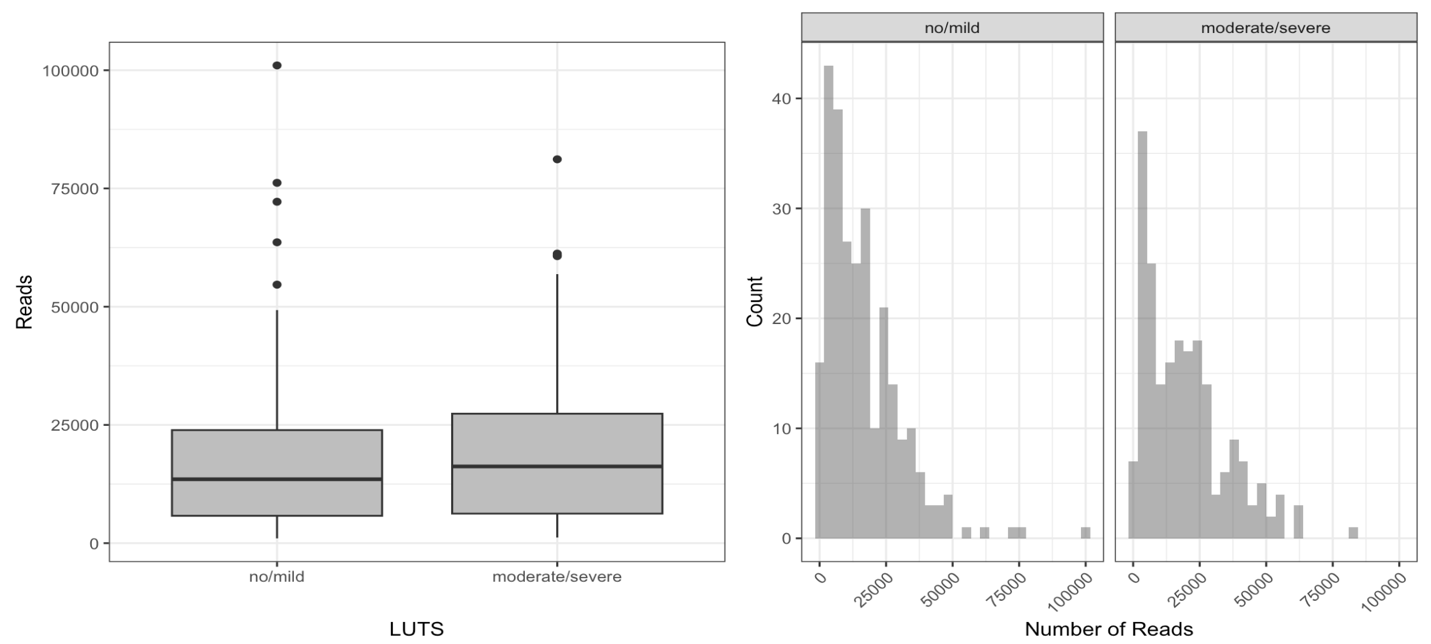

**Figure S5**. Boxplot and histogram of the number of reads by LUTS severity. No/mild LUTS (red) has significantly less reads than moderate/severe LUTS (blue, p = 0.04).

**Table S1.** Associations of clinical characteristics and urobiome alpha diversity. Categorical data are reported as median [interquartile range (IQR)] of the diversity measure for the specified group, continuous variables (age) are reported as correlation coefficients. Significant associations were identified between BPH and Shannon, Inverse Simpson, and Pielou indices are significant (p < 0.05, bolded) determined by Wilcoxon rank sum test.

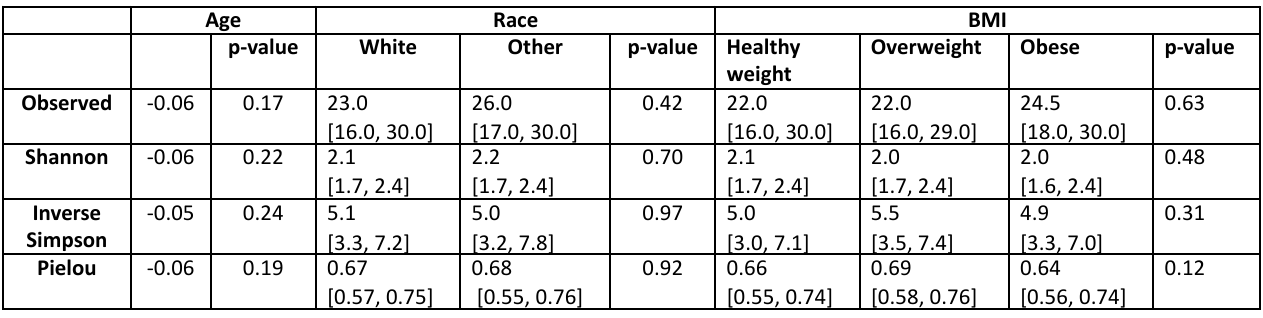

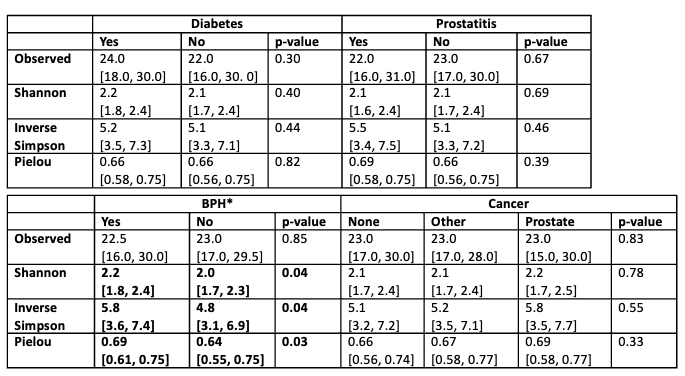

**Table S2**. Overall LUTS and obstructive and irritative symptom scores summarized for each urotype. There were no significant differences in the number of men with moderate to severe LUTS in each cluster, nor in the obstructive and irritative symptom scores.

| **Urotype** | **LUTS score (%),**  **Moderate/severe** | **Score (median, [IQR])**  **Obstructive Irritative** | |
| --- | --- | --- | --- |
| **1 (n=85)** | 35 (41.2%) | 1.0 [0.0, 5.0] | 4.0 [2.0, 6.0] |
| **2 (n=89)** | 43 (48.3%) | 3.0 [1.0, 7.0] | 4.0 [3.0, 7.0] |
| **3 (n=65)** | 27 (41.5%) | 1.0 [0.0, 4.0] | 4.0 [3.0, 7.0] |
| **4 (n=61)** | 24 (39.3%) | 3.0 [0.0, 5.0] | 3.0 [2.0, 6.0] |
| **5 (n=59)** | 33 (55.9%) | 3.0 [1.0, 6.0] | 5.0 [3.0, 7.0] |
| **6 (n=58)** | 27 (46.6%) | 3.0 [0.0, 5.8] | 3.0 [2.0, 6.0] |
| **7 (n=37)** | 16 (43.2%) | 2.0 [0.0, 5.0] | 4.0 [2.0, 6.0] |
| **8 (n=19)** | 4 (21.1%) | 2.0 [0.0, 4.5] | 3.0 [2.0, 5.0] |
| **p-value** | 0.23 | 0.10 | 0.31 |

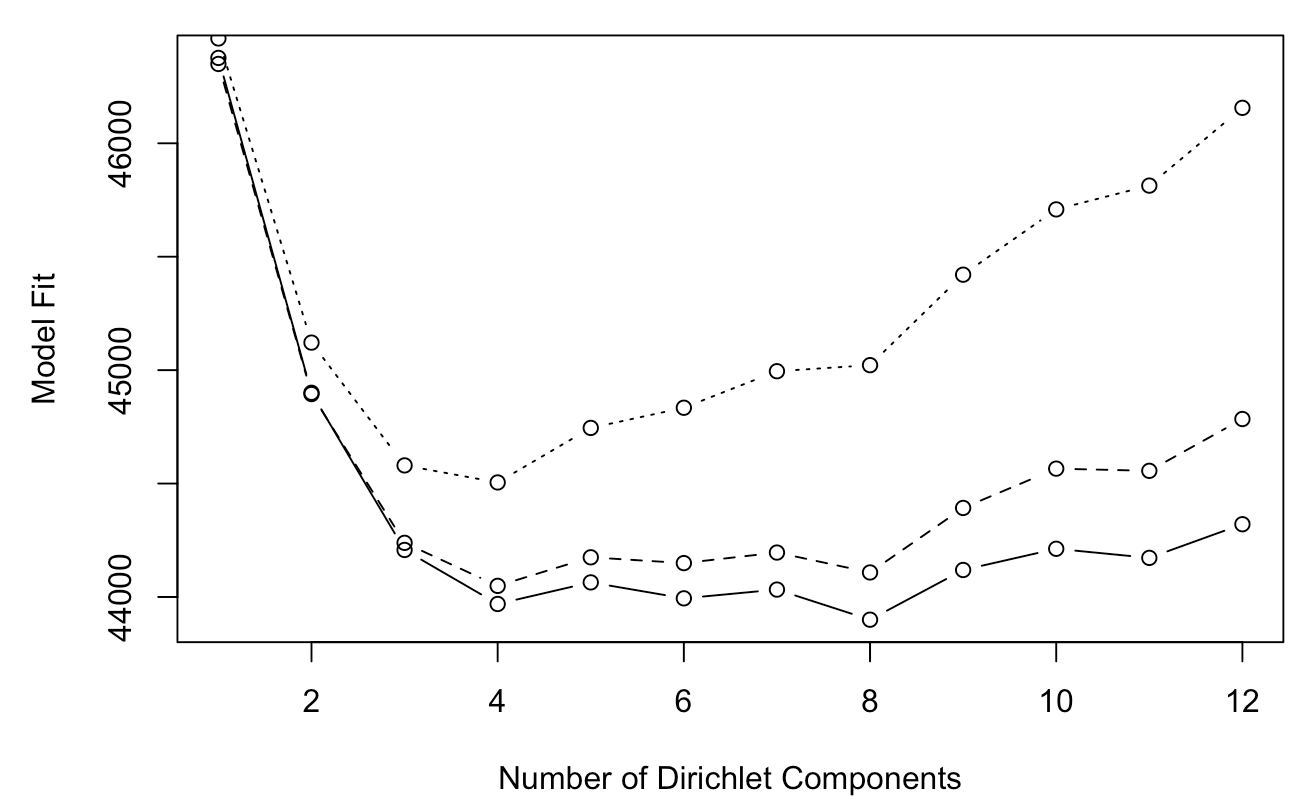

**Figure S5**. Evaluation of the DMM model fit of varying numbers of clusters using the Laplace approximation, Aikaike Information Criterion (AIC), and Bayesian Information Criterion (BIC). Smallest dotted line represents AIC, middle line is the BIC, and solid line is the Laplace approximation.
